## Supplementary figures and images for "Tetraspanin Cd9b and Cxcl12a/Cxcr4b have a synergistic effect on the control of collective cell migration"

### Fig_S1.tif

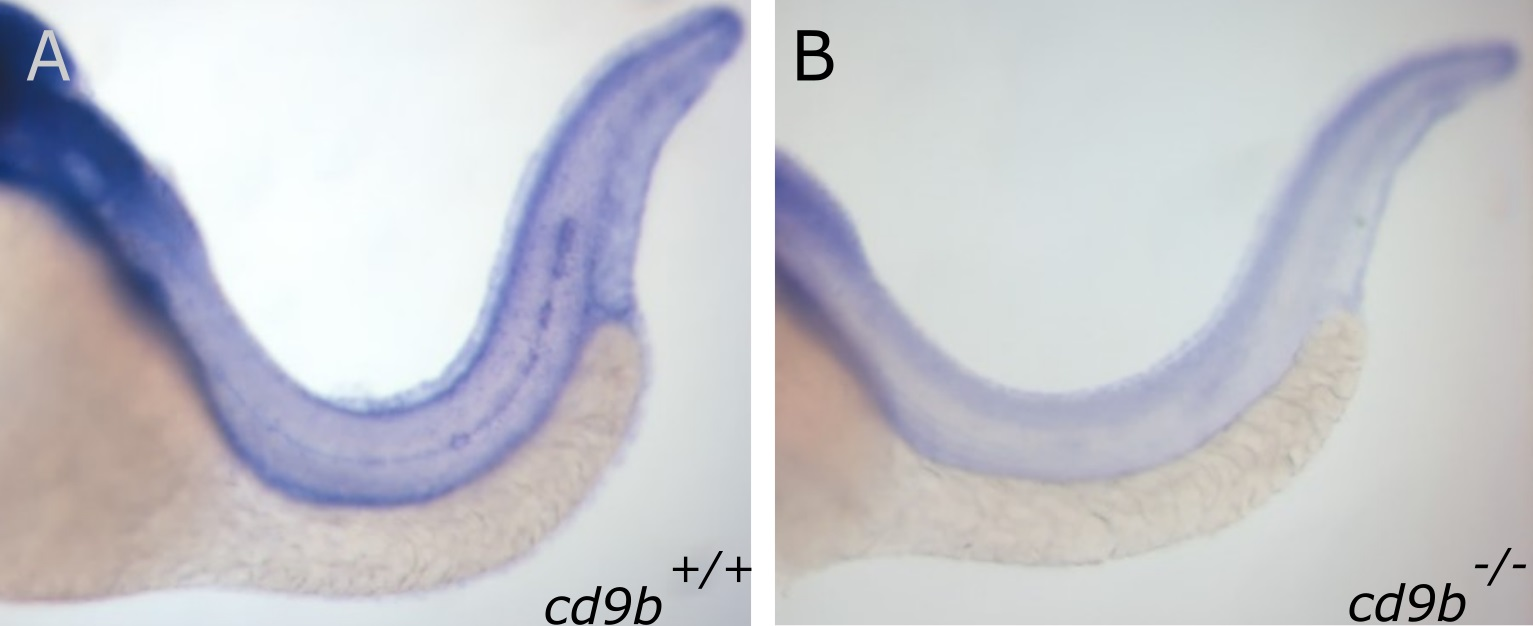

### Fig_S2.tif

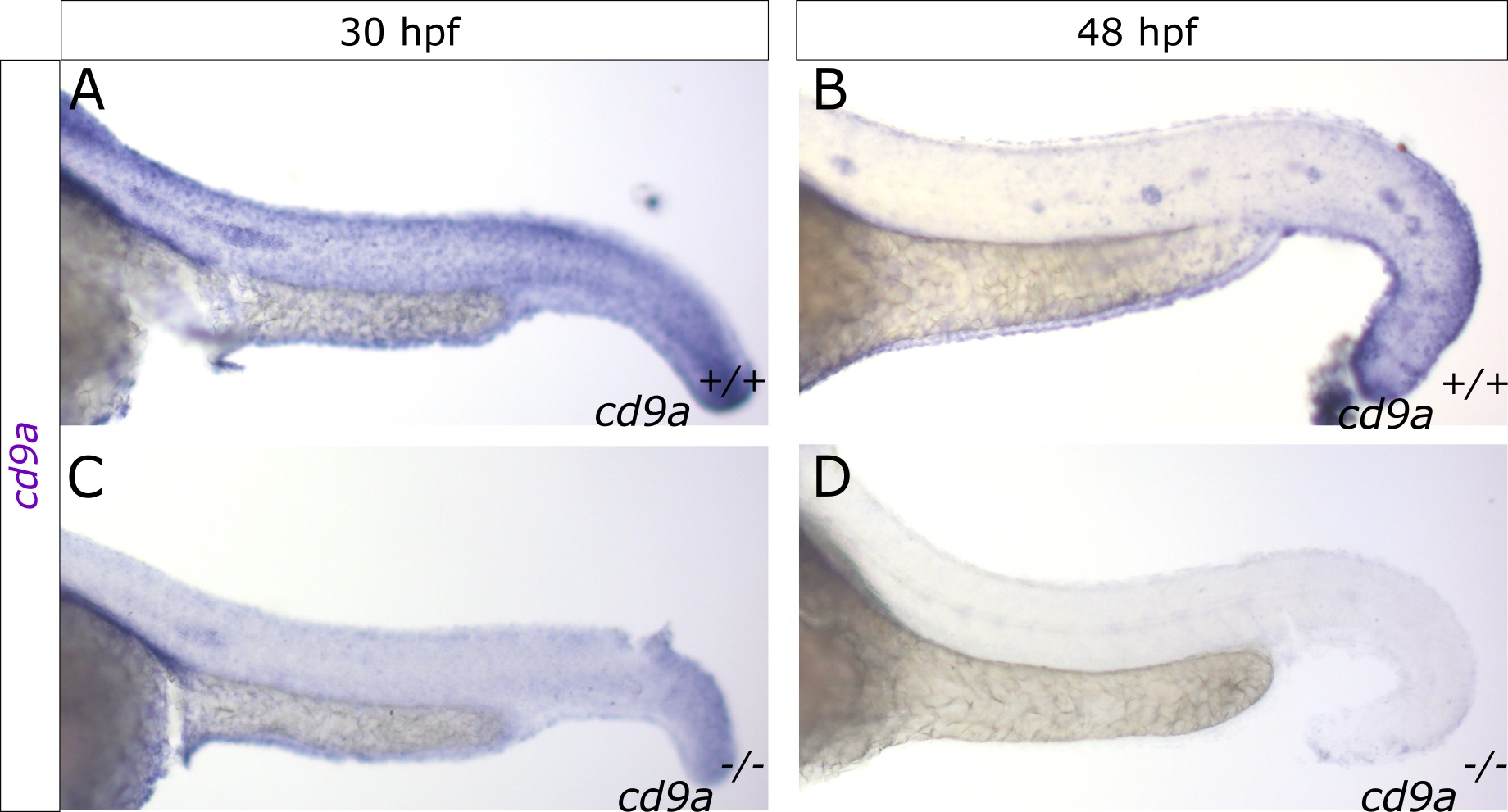

### Fig_S3.tif

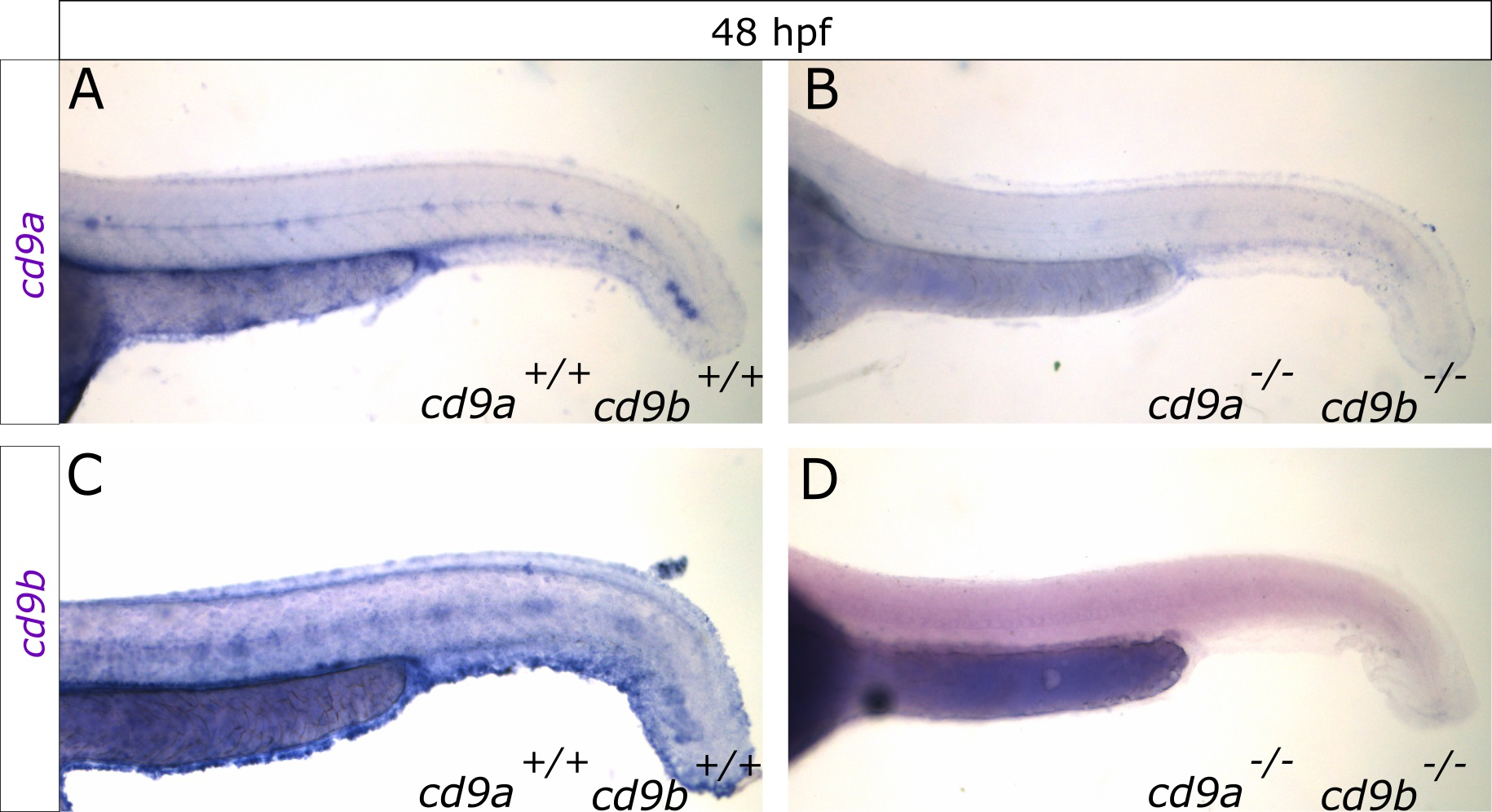

### Fig_S4.tif

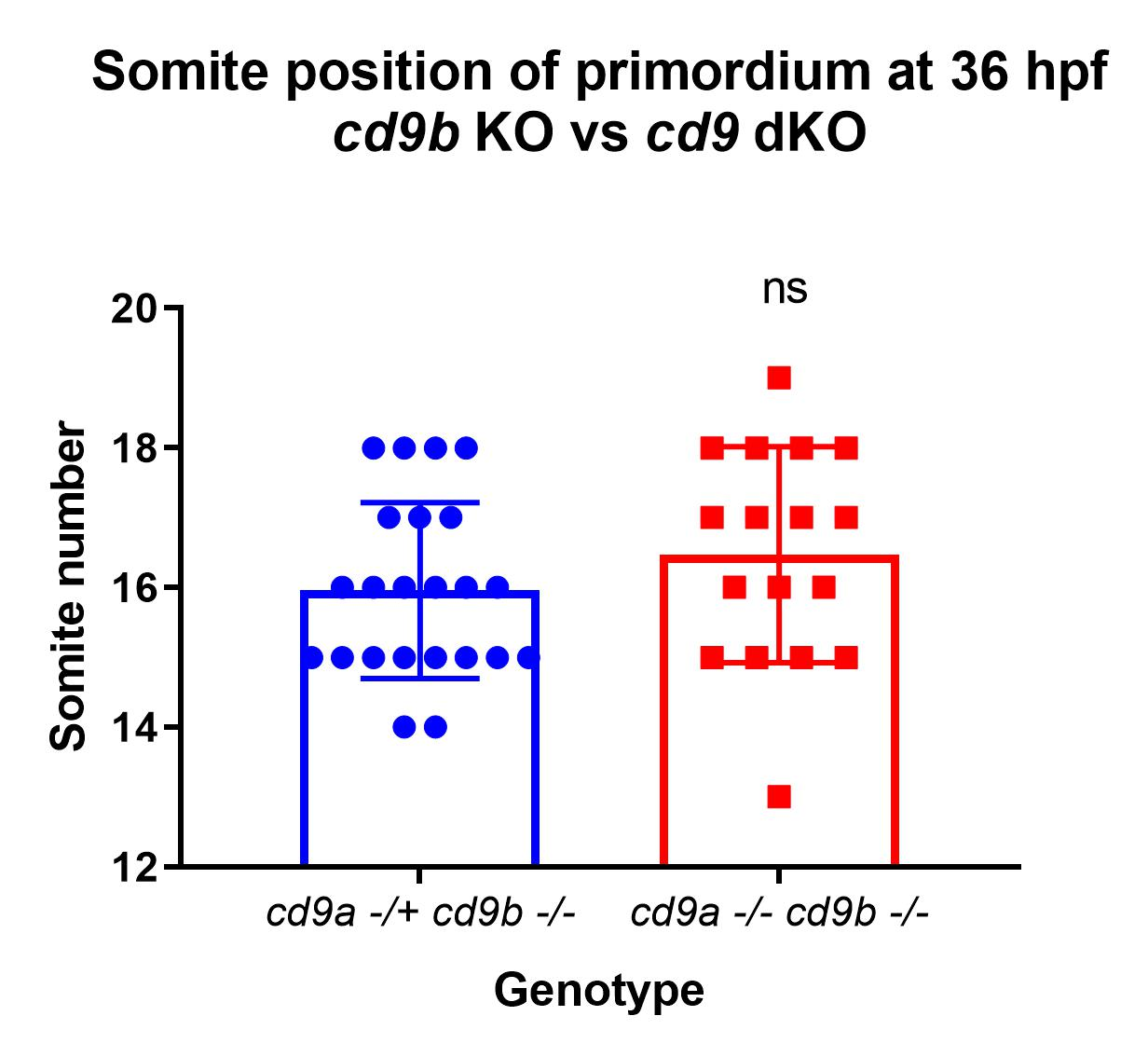

### Fig_S5.tif

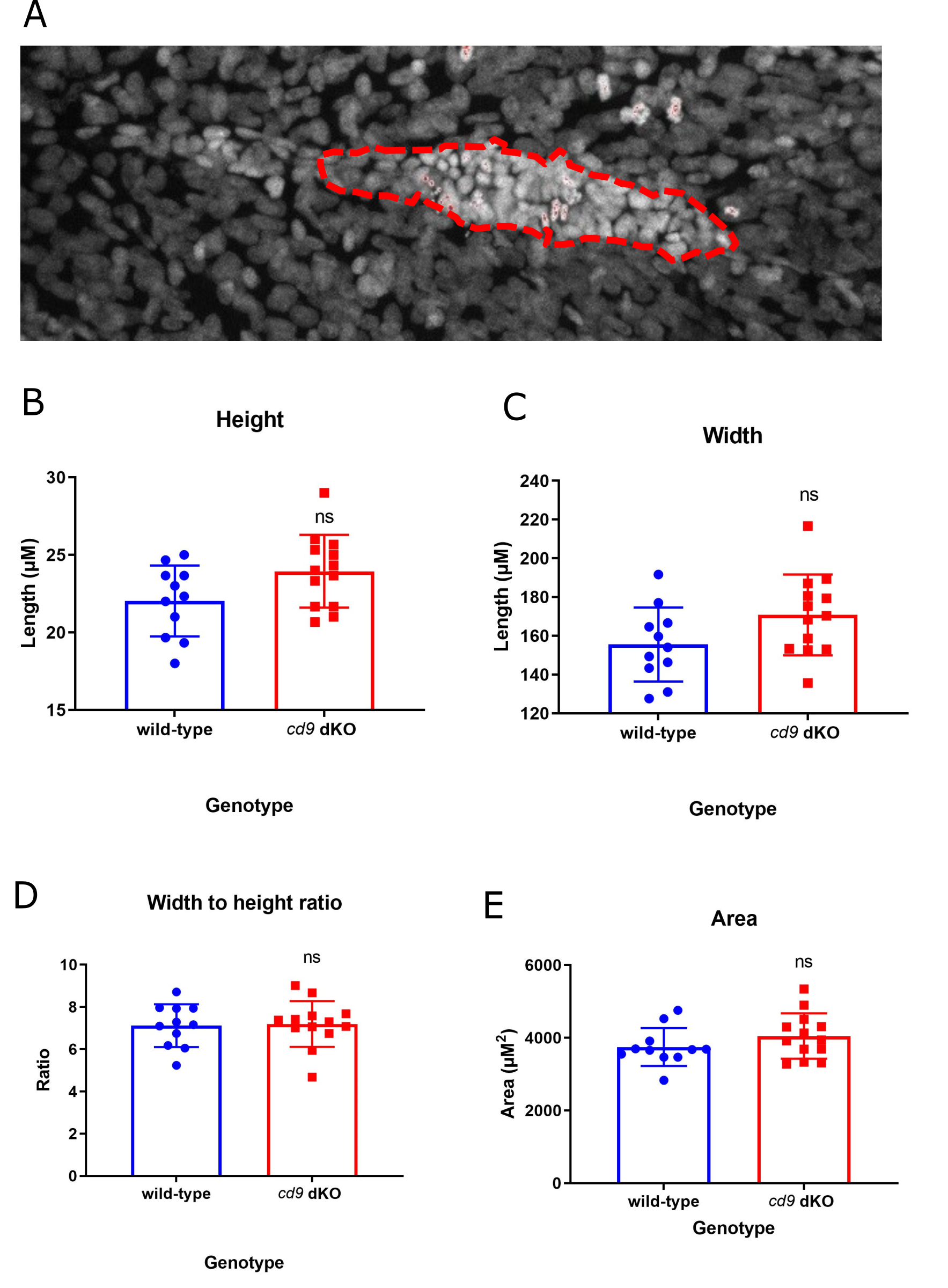

### Fig_S6.tif

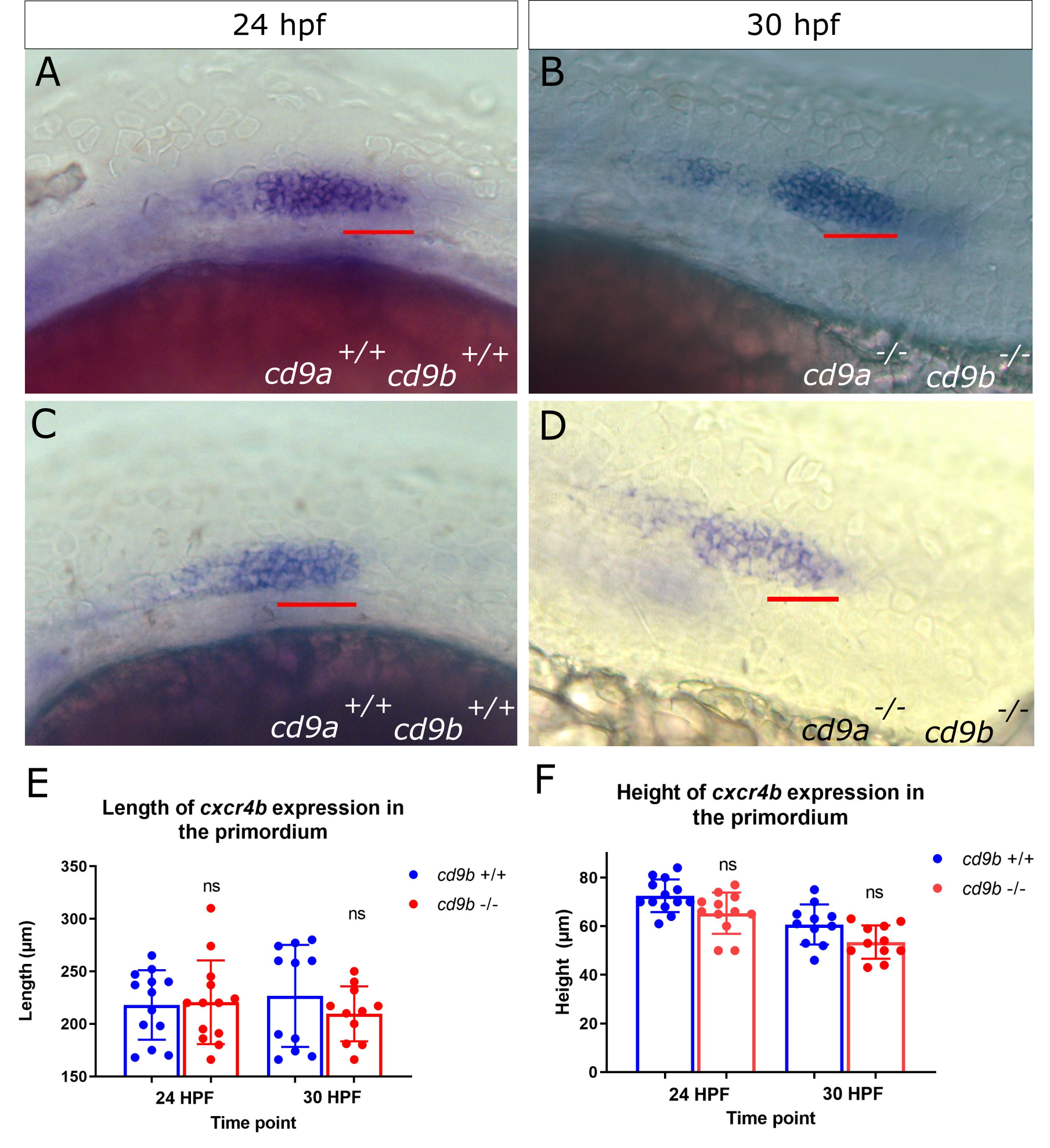

### Fig_S7.tif

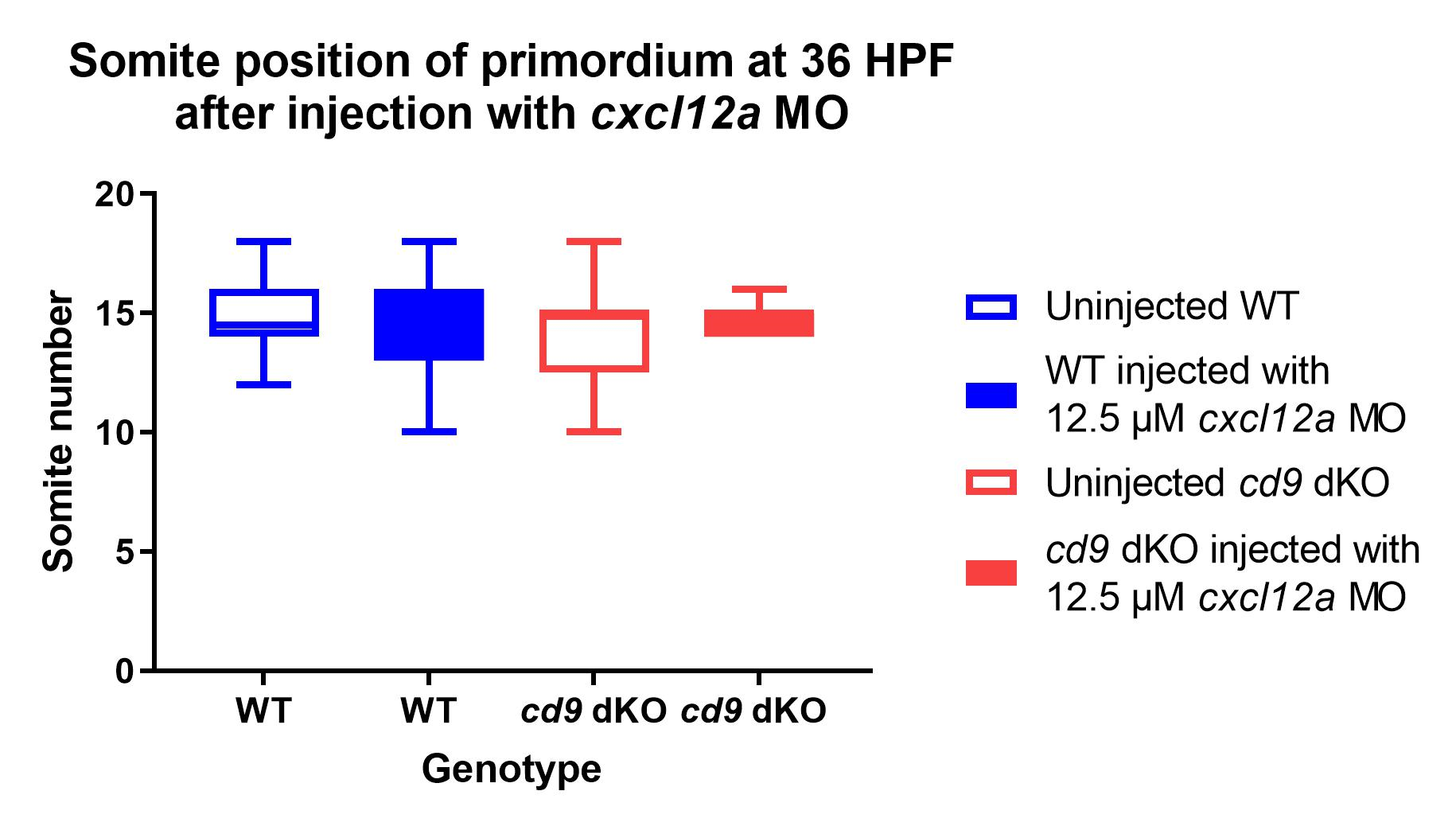
